# Statistical prediction of the future impairs episodic encoding of the present

**DOI:** 10.1101/851147

**Authors:** Brynn E. Sherman, Nicholas B. Turk-Browne

## Abstract

Memory is typically thought of as enabling reminiscence about past experiences. However, memory also informs and guides processing of future experiences. These two functions of memory are often at odds: remembering specific experiences from the past requires storing idiosyncratic properties that define particular moments in space and time, but by definition such properties will not be shared with similar situations in the future and thus may not be applicable to future situations. We discovered that, when faced with this conflict, the brain prioritizes prediction over encoding. Behavioral tests of recognition and source recall showed that items allowing for prediction of what will appear next based on learned regularities were less likely to be encoded into memory. Brain imaging revealed that the hippocampus was responsible for this interference between statistical learning and episodic memory. The more that the hippocampus predicted the category of an upcoming item, the worse the current item was encoded. This competition may serve an adaptive purpose, focusing encoding on experiences for which we do not yet have a predictive model.

## Introduction

Human memory contains two fundamentally different kinds of information — episodic and statistical. Episodic memory refers to the encoding of specific details of individual experiences (e.g., what happened on your last birthday), whereas statistical learning refers to extracting what is common across multiple experiences (e.g., what tends to happen at birthday parties). Episodic memory allows for vivid recollection and nostalgia about past events, whereas statistical learning leads to more generalized knowledge that affords predictions about new situations. Episodic memory occurs rapidly and stores even related experiences distinctly in order to minimize interference, whereas statistical learning occurs more slowly and overlays memories in order to represent their common elements or regularities. Given these behavioral and computational differences, theories of memory have argued that these two kinds of information must be processed serially and stored separately in the brain (McClelland et al., 1995; Squire, 2004): episodic memories are formed first in the hippocampus and these memories in turn provide the input for later statistical learning in the neocortex as a result of consolidation (Frankland and Bontempi, 2005; Richards et al., 2014; Tompary and Davachi, 2017).

Here we reveal a relationship between episodic memory and statistical learning in the reverse direction, whereby learned regularities determine which memories are formed in the first place. Specifically, we examine whether the ability to predict what will appear next — a signature of statistical learning — reduces encoding of the current experience into episodic memory. This hypothesis depends on two theoretical commitments: first, that the adaptive function of memory is to guide future behavior by generating expectations based on prior experience (Schacter et al., 2017); second, that memory resources are limited, because of attentional bottlenecks that constrain encoding (Aly and Turk-Browne, 2017) and/or because new encoding interferes with the storage or retrieval of existing memories (Shiffrin and Atkinson, 1969). Accordingly, in allocating memory resources, we propose that it is less important to encode an ongoing experience when it already generates strong expectations about future states of the world. When the current experience affords no such expectations, however, encoding it into memory provides the opportunity to extract new, unknown regularities that enable more accurate predictions in subsequent encounters. After demonstrating this role for statistical learning in episodic memory behaviorally, we identify an underlying mechanism in the brain using fMRI, based on the recent discovery that both processes depend upon the hippocampus and thus compete to determine its representations and output (Schapiro et al., 2017).

## Results

### Experiment 1a

We exposed human participants to a stream of pictures and later tested their memory (Figure 1A). The pictures consisted of outdoor scenes from 12 different categories (e.g., beach, mountain, farm). Three of the categories (type A, predictive) were each reliably followed by one of three other categories (type B, predictable); the remaining six categories (type X, non-predictive, non-predictable) were randomly inserted into the stream. That is, every time participants saw a picture from an A category, they always saw a picture from a specific B category next; however, when a picture from an X category appeared, it was variably preceded and followed by pictures from several other categories (Figure 1B). Participants were not informed about these predictive A → B category relationships and learned them incidentally through exposure (Brady and Oliva, 2008). Although each category was shown several times, every individual picture in the stream was a novel exemplar from the category and shown only once. For example, whenever a picture from the beach category appeared, it was a new beach that they had not seen before. After the stream, we tested memory for these individual pictures amongst new exemplars from the same categories. The key hypothesis was that exemplars from predictive categories would be remembered worse than exemplars from non-predictive categories.

**Figure 1:**
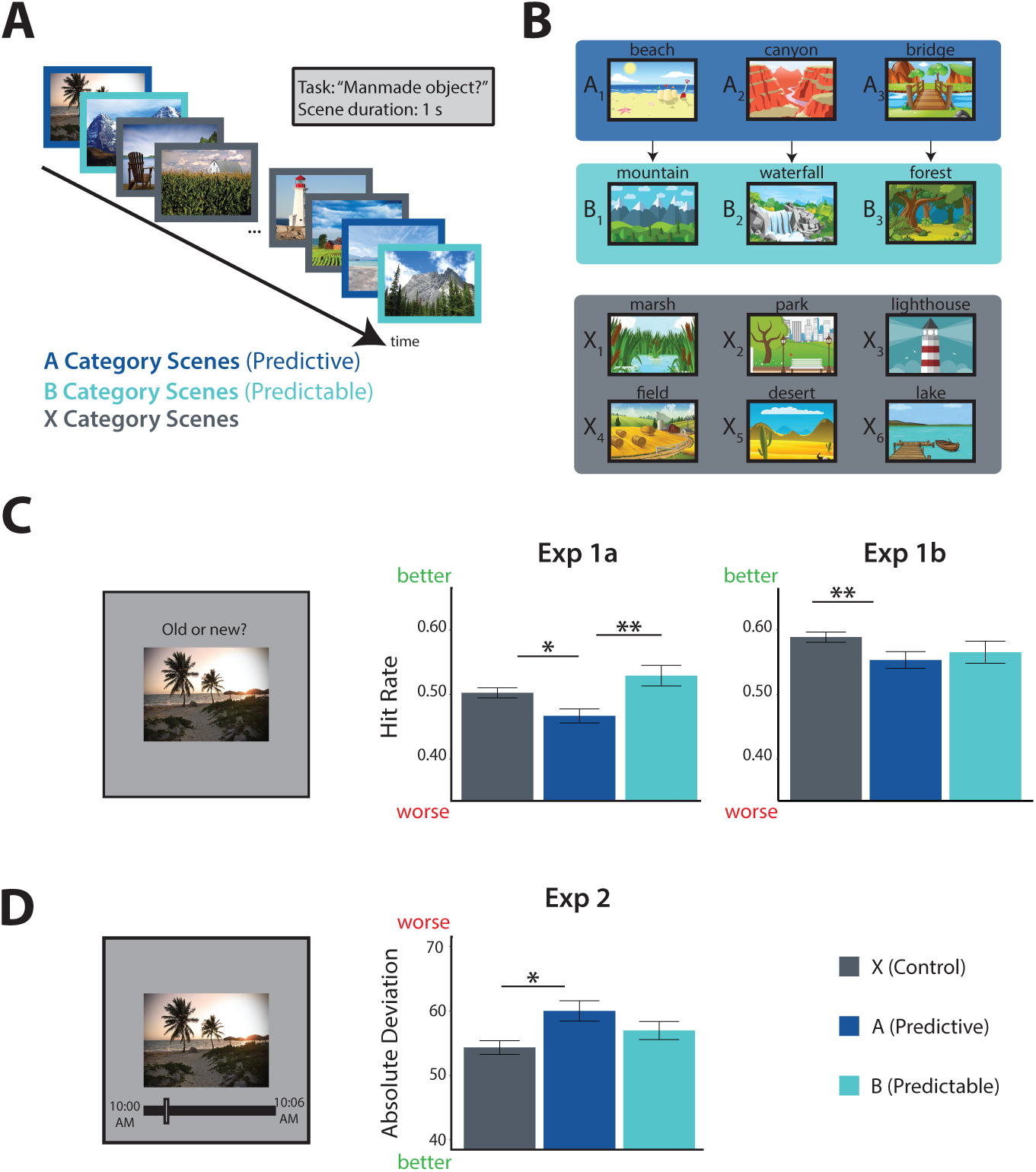
Behavioral Experiments. A) Task design: participants viewed a continuous stream of scene pictures, during which they made a judgment of whether or not there was a manmade object in the scene. B) Example scene category pairings for one participant: 3 of 12 categories were assigned to condition A; each was reliably followed by one of 3 different categories assigned to condition B (illustrated by arrows). The remaining 6 categories were assigned to condition X and were not consistently preceded or followed by any particular category. C) Left: surprise recognition memory test. Middle: proportion of old exemplars recognized (hit rate) as a function of condition (higher hit rate is better memory) for Experiment 1a. Right: hit rate for Experiment 1b. D) Left: temporal source memory test. Right: absolute difference between reported and actual time of encoding as a function of condition (higher deviation is worse memory). Error bars reflect within-participant standard error of the mean. *p *<*0.05,**p *<*0.01.

### Encoding Phase

While viewing the stream, 30 participants performed a cover task in which they judged whether or not there was a manmade object in the scene. Participants performed quite well on this task (mean accuracy = 0.91). This performance level was reliably above chance (0.5; t(29) = 42.38, p *<*0.001). Assessing response times over the course of the experiment, we found a reliable main effect of experiment quartile (F(3,87) = 8.30, p *<*0.001), a marginal main effect of condition (F(2,58) = 3.09, p = 0.053), and a marginal interaction between condition and quartile (F(6,174) = 2.15, p = 0.050). This interaction reflected growing facilitation for the predictable B category, with marginally faster response times by the fourth quartile relative to the X (t(29) = 2.02, p = 0.053) and A (t(29) = 1.99, p = 0.056) categories, whose appearance could not be predicted (Hunt and Aslin, 2001; Olson and Chun, 2001).

### Test Phase

To evaluate overall episodic memory performance, we calculated A′ for each participant as a non-parametric measure of sensitivity. All participants had memory performance numerically above the chance level of 0.5 (mean A*!* = 0.72, t(29) = 20.02, p *<*0.001; mean hit rate = 0.50; mean false alarm rate = 0.23). We did not find a reliable main effect of condition on A*!* (F(2,58) = 2.37, p = 0.10). However, A′ takes into account both the hit rate and the false alar rate. Given our hypothesis of worse *encoding* for the old exemplars from the predictive A categories, we expected that hit rate would be a more sensitive measure.

Indeed, there was a main effect of condition on hit rate (F(2,58) = 4.75, p = 0.012), with a lower hit rate for pictures from the A categories relative to both B (t(29) = -2.79, p = 0.0092) and X (t(29) = -2.33, p = 0.027) categories (Figure 1C, middle. There was no difference in hit rate between B and X categories (t(29) = 1.19, p = 0.24), showing that the memory deficit is selective to whether a category was predictive (A vs. X), not whether it was predictable (B vs. X). As hypothesized, this memory deficit reflected a failure to encode specific A exemplars rather than a generic impairment for A categories (De Brigard et al., 2017; Smith et al., 2013), as the false alarm rate for new exemplars from each category at test did not differ by condition (F(2, 58) = 0.29, p = 0.75). Given these findings, analyses in subsequent experiments consider hit rate and false alarm rate separately by condition.

If prediction from statistical learning interferes with episodic memory encoding, we might expect the memory deficit for predictive A items to increase as learning progresses. We thus analyzed hit rate across conditions as a function of the encoding phase quartile in which a picture was presented (Figure S1A). We found main effects of both quartile (F(3, 87) = 3.49, p = 0.019) and condition (F(2, 58) = 4.75, p = 0.012), but no interaction (F(6, 174) = 0.74, p = 0.62). Focusing later within the encoding phase, we found reliable main effects of condition in the third (F(2, 58) = 4.09, p = 0.022) and fourth quartiles (F(2,58) = 4.12, p = 0.021). In the third quartile, memory for A was reliably worse than for B (t(29) = -2.73, p = 0.011), but not X (t(29) = -0.40, p = 0.69). In the fourth quartile, memory for A was reliably worse than for both B (t(29) = -2.55, p = 0.016) and X (t(29) = -2.48, p = 0.019). In contrast, we did not find any reliable main effects or pairwise differences between conditions in the first or second quartiles. These results are consistent with the deficit for predictive A items emerging over time during statistical learning.

### Experiment 1b

Experiment 1a suggested that prediction from statistical learning can impair episodic memory. Most critically, A items that were predictive of an upcoming B item were remembered worse than non-predictive X items. To establish the robustness of these results, here we performed a pre-registered and highly powered online replication study.

### Encoding Phase

A group of 64 participants on the online data collection platform Prolific participated in the same encoding task as Experiment 1a. Participants performed quite well on the cover task (mean accuracy = 0.92; relative to 0.5 chance: t(63) = 67.32, p *<*0.001). Examining response times over the course of learning, we again found a reliable main effect of quartile (F(3,189) = 15.13, p *<*0.001), but no main effect of condition (F(2,126) = 0.018, p = 0.98), nor an interaction between condition and quartile (F(6,378) = 0.73, p = 0.63).

### Test Phase

To examine overall memory performance we again measured A′ for each participant and verified that memory performance at the group level was above chance (mean |A ′ = 0.69, t(63) = 19.77, p *<*0.001; mean hit rate = 0.57; mean false alarm rate = 0.35).

As in Experiment 1a, we did not find a reliable main effect of condition on A′ (F(2,126) = 1.59, p = 0.21) or false alarm rate (F(2,126) = 0.060, p = 0.94). However, we did find a marginal main effect of condition on hit rate (F(2,126) = 2.76, p = 0.067; Figure 1C, right). Critically, we robustly replicated the key pairwise difference in hit rate between A and X categories (t(63) = -3.03, p = 0.0036), the comparison that isolates the effect of predictiveness. Unlike Experiment 1a, the difference in hit rate between A and B categories was not reliable (t(63) = -0.68, p = 0.50), though memory for A was still numerically lower than B. The B and X categories again did not differ in hit rate (t(63) = -1.46, p = 0.15).

We again considered whether the pattern of results changed over the course of learning (Figure S1B). Assessing hit rate as a function of encoding quartile, we found a reliable main effect of quartile (F(3, 189) = 4.95, p = 0.0025), a marginal main effect of condition (F(2, 126) = 2.76, p = 0.067) and a marginal interaction (F(6, 378) = 1.79, p = 0.099). The main effect of condition was reliable only within the fourth quartile (F(2, 126) = 5.60, p = 0.0047), with memory for A reliably worse than for both B (t(63) = -2.20, p = 0.031) and X (t(63) = -3.67, p *<*0.001).

### Experiment 2

Together, Experiments 1a and 1b demonstrated that episodic encoding is worse for predictive vs. non-predictive pictures using a surprise recognition memory test. We interpret this result as evidence of competition between prediction and encoding in the hippocampus. However, recognition tests do not definitively probe aspects of episodic memory that specifically depend on the hippocampus. Participants could have relied upon a generic sense of familiarity with the pictures, which can be supported by cortical areas (Brown and Aggleton, 2001; Davachi et al., 2003; Norman and O’Reilly, 2003). We thus designed Experiment 2 with a different, recall-based memory test. After encoding the same kind of picture stream, participants were unexpectedly asked at test to indicate at what exact time (on the clock) they had seen each picture in the stream. As before, encoding of the time was incidental as they were not informed in advance that they would be tested. This kind of precise temporal source memory requires the retrieval of details about the context in which each picture was encoded, a hallmark function of episodic memory (e.g., remembering who arrived first at a birthday party) that critically depends upon the hippocampus (Davachi and DuBrow, 2015; Miller et al., 2013; Mitchell and Johnson, 2009).

### Encoding Phase

A group of 30 new participants performed quite well on the same manmade cover task as Experiments 1a (mean accuracy = 0.93; relative to 0.5 chance: t(29) = 44.07, p *<*0.001). There was again a main effect of experiment quartile on response times (F(3,87) = 7.82, p *<*0.001), but no main effect of condition (F(2,58) = 0.22, p = 0.80) nor an interaction between condition and quartile (F(6,174) = 0.69, p = 0.65).

### Test Phase

In the memory test, participants were presented with a picture and first asked to indicate whether they thought it was old or new, i.e., whether it was presented during the encoding stream. If they indicated that the picture was old, they were then asked to recall when during the stream they had seen the picture. We included the initial old/new recognition judgment because we felt that it would be awkward to ask participants to report the time of a picture they did not remember seeing previously. Nevertheless, our primary hypothesis was that precision of temporal source memory recall would be lower for exemplars from predictive categories, resulting in greater deviation or error for A compared to X categories.

In terms of overall recognition memory from the initial old/new judgments, all participants had an A′ numerically above the chance level of 0.5 (mean A′ = 0.75, t(29) = 23.31, p *<*0.001; mean hit rate = 0.38; mean false alarm rate = 0.11). Neither the hit rate (F(2,58) = 0.74, p = 0.48) nor the false alarm rate (F(2,58) = 0.33, p = 0.72) differed by condition. The lack of a hit rate effect differed from Experiment 1, but we suspect that this may be an artifact of introducing the source memory task. Specifically, participants in Experiment 2 knew that responding “old” would prompt a difficult follow-up question about their temporal source memory. As a result, they may have strategically adopted a more conservative criterion to avoid the source judgment unless they had a strong memory with high confidence. Consistent with this interpretation, participants were less likely to respond “old” in general in Experiment 2 (mean proportion “old” responses = 0.33) than in Experiment 1a (0.45; t(58) = 4.76, p *<*0.001).

Regardless, our focus in this experiment was on temporal source memory recall. We assessed overall source memory by computing the average absolute deviation from the correct clock time for all hits. Higher absolute deviation indicates *lower* precision in memory. The mean absolute deviation across participants was 56.3 pictures, or 84.5 seconds. Twenty-six of the 30 participants had a mean deviation numerically lower than chance (determined via permutation test to be 63.8 pictures), and thus performance at the group level was reliably above chance (t(29) = -7.04, p *<*0.001). As hypothesized, there was a main effect of condition on absolute deviation (Figure 1D; F(2,58) = 3.17, p = 0.049). Pictures from the A categories had greater deviation (less precision) than those from the X categories (t(29) = 2.26, p = 0.031), which differed only in that they were not predictive of the upcoming category. Precision was also lower for A relative to B categories, but this difference did not reach significance (t(29) = 1.45, p = 0.16); B and X categories also did not reliably differ from each other (t(29) = 1.23, p = 0.23).

Examining how memory changed over the course of learning, as we did in the preceding experiments, is difficult here because of the use of a temporal source memory measure. Namely, because the range of possible deviations changes over time (i.e., by chance, pictures from the middle of encoding would have lower average deviation than from the beginning and end), examining how source memory changes over time is confounded. Moreover, because of the lower hit rate in this experiment, there are fewer responses per quartile and in fact three participants did not have any responses in at least one quartile. We performed the analysis nevertheless, for completeness, and found a main effect of quartile (F(3, 78) = 41.9, p *<*0.001), a marginal main effect of condition (F(2, 52) = 2.73, p = 0.075), but no interaction (F(6, 156) = 1.5, p = 0.18). Follow-up tests did not reveal any reliable effects within individual quartiles.

### Experiment 3 (fMRI)

Across Experiments 1a, 1b, and 2, we found robust and consistent evidence that memory for predictive A items is reduced relative to the non-predictive, non-predictable control X items. What explains this worse encoding of predictive pictures? We propose that this results from the co-dependence of statistical learning and episodic memory on the hippocampus (Schapiro et al., 2017). Specifically, we hypothesize that the appearance of a picture from an A category triggers the retrieval and predictive representation of the corresponding B category in the hippocampus. This in turn prevents the hippocampus from encoding a new representation of the specific details of that particular A picture, which would be needed for later recall from episodic memory. To evaluate this hypothesis, Experiment 3 employed high-resolution fMRI during the encoding phase to link hippocampal prediction to subsequent memory.

### Encoding Phase Behavior

A group of 36 new participants performed the same manmade cover task as the Experiments 1a, 1b, and 2. Performance across all runs (including the templating phases; see Materials and Methods) remained quite high (mean accuracy = 0.94; relative to 0.5 chance: t(35) = 18.75, p *<*0.001). Response times were examined across thirds of the encoding phase rather than quartiles because there were three fMRI runs in this phase. As in Experiment 1a, we found a pattern of growing facilitation for the B categories. Although there were no main effects of experiment third (F(2,68) = 0.44, p = 0.65) or condition (F(2,68) = 1.24, p = 0.29), nor an interaction (F(4,136) = 1.17, p = 0.33), response times in the third run were reliably faster for B categories relative to X categories (t(34) = 2.23, p = 0.033); the difference for B relative to A categories was in the same direction but not reliable (t(34) = 1.39, p = 0.17).

### Test Phase Behavior

In Experiment 3, we returned to the recognition memory task from Experiments 1a and 1b. All participants exhibited A′ above the chance level of 0.5 except for one, who was excluded from all other behavioral and fMRI analyses (all participants: mean A′ = 0.68, t(36) = 14.63, p *<*0.001; mean hit rate = 0.61; mean false alarm rate = 0.39). Neither hit rate (F(2,70) = 2.23, p = 0.12) nor false alarm rate (F(2,70) = 0.83, p = 0.44) differed by condition. Notably, this experiment differed from the behavioral experiments in that participants first completed a “pre” templating phase (necessary for the fMRI decoding analyses; see Materials and Methods), in which they viewed pictures from scene categories — that would subsequently be paired in the encoding phase — in a random order. Exposure to randomness prior to structure can impede statistical learning (Jungé et al., 2007), which might help to explain the diminished memory effect in this experiment. This is consistent with weakened behavioral evidence of statistical learning in other multivariate fMRI studies with templating phases (Schapiro et al., 2012). More generally, it is not uncommon for behavioral effects to be smaller in fMRI studies, including in prior statistical learning studies (Turk-Browne et al., 2009). Although we did not observe an overall memory effect at the group level, we followed precedent (Wimmer and Shohamy, 2012; Schlichting et al., 2015) in leveraging individual differences in learning to examine the relationship between memory behavior and neural measures across participants.

### Neural Decoding of Perceived Information

The primary purpose of the fMRI experiment was to measure neural prediction during statistical learning in the encoding phase. We used a multivariate pattern classification approach (Cohen et al., 2017), which quantified neural prediction of B categories during the encoding of A pictures. Classification models were trained for each category based on patterns of fMRI activity in a separate phase of the experiment (“pre” templating phase; see Materials and Methods), during which participants were shown pictures from all categories in a random order. These classifiers were then tested during viewing of the encoding stream containing category pairs, providing a continuous readout of neural evidence for each category. We performed this analysis based on fMRI activity patterns from the hippocampus, our primary region of the interest (ROI), as well as from control ROIs in occipital and parahippocampal cortices. These control ROIs were chosen because as visual areas we expected them to be sensitive to the category of the current picture being viewed but not necessarily to predict the upcoming B category given an A picture.

To validate this approach, we first trained and tested classifiers on the viewing of pictures from the A categories (“Perception of A”) and B categories (“Perception of B”) (Figure 2). A categories could be reliably decoded in the occipital (t(35) = 4.34, p *<*0.001) and parahippocampal (t(35) = 3.83, p *<*0.001) cortices, but not in the hippocampus (t(35) = -0.17, p = 0.87; main effect of region: F(2,70) = 7.47, p = 0.0012). In contrast, B categories could be reliably decoded in all three regions (occipital: t(35) = 5.96, p *<*0.001, parahippocampal: t(35) = 3.52, p = 0.0012, hippocampus: t(35) = 2.26, p = 0.030; main effect of region: F(2,70) = 6.73, p = 0.0021). Combining both perception conditions, there was a reliable main effect of region (F(2,70) = 14.89, p *<*0.001), but no main effect of condition (F(1,35) = 2.06, p = 0.16) and no interaction between region and condition (F(2,70) = 0.94, p = 0.39).

**Figure 2:**
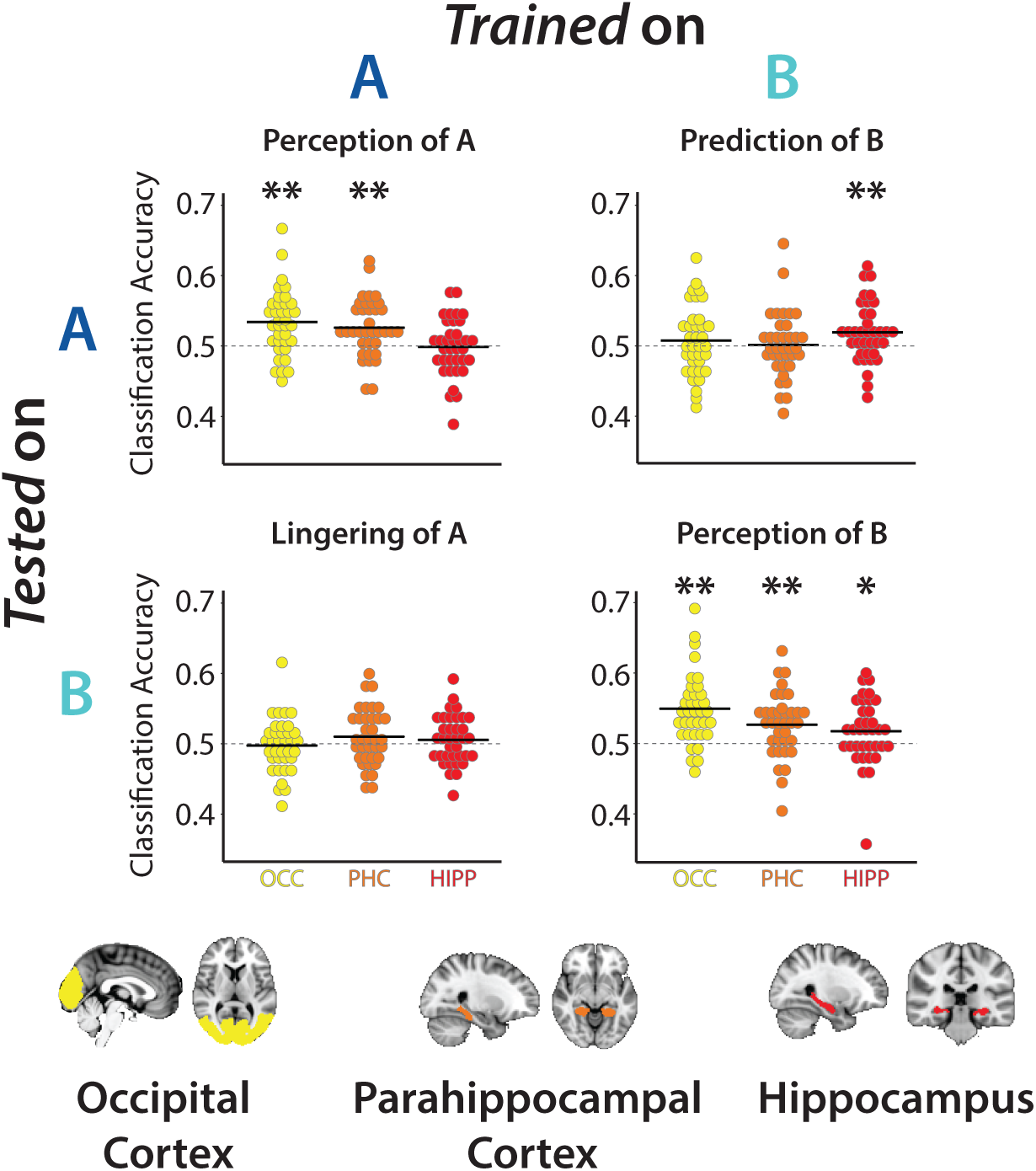
Category Decoding in fMRI Experiment. Top: Classification accuracy in occipital cortex (OCC), parahippocampal cortex (PHC), and hippocampus (HIPP) for each of the four combinations of training or testing on A or B categories. For every A/B combination and ROI, each dot is one participant and the black line is the mean across participants. Bottom: Regions of interest. HIPP and PHC were manually segmented in native participant space (transformed into standard space for visualization); OCC was defined in standard space and transformed into native participant space.

The hippocampus was unique in showing reliable decoding during the Perception of B but not Perception of A. Although this difference did not reach not significance (t(35) = -1.63, p = 0.11), we sought to determine whether it reflected a diminished representation of A or an enhanced representation of B. To establish a baseline, we trained and tested classifiers on the viewing of pictures from the control X categories (Figure S2). Mirroring the Perception of A results, X categories could be reliably decoded in the occipital (t(35) = 7.16, p *<*0.001) and parahippocampal (t(35) = 2.47, p = 0.019) cortices, but not in the hippocampus (t(35) = -0.49, p = 0.63; main effect of region: F(2,70) = 34.61, p *<*0.001). Within the hippocampus, we found a marginal main effect of condition (F(2,70) = 2.58, p = 0.083), with X lower than B (t(35) = -2.14, p = 0.039) but not A (t(35) = -0.10, p = 0.92). These results suggest that prediction may enhance the representation of predictable items in the hippocampus.

### Neural Decoding of Predicted Information

We next tested the hypothesis that the hippocampus predicts B categories during viewing of the associated A categories. We trained classifiers on pictures from each B category and tested on pictures from the corresponding A category (“Prediction of B”). Crucially, the upcoming B category could be decoded during A in the hippocampus (t(35) = 2.73, p = 0.0098), but this was not possible in occipital (t(35) = 0.94, p = 0.35) or parahippocampal (t(35) = 0.17, p = 0.87) cortices (main effect of region: F(2,70) = 1.76, p = 0.18). Control analyses ruled out potential confounds related to the timing of the fMRI signal: training classifiers on A and testing on B categories (“Lingering of A”), did not yield reliable decoding in the hippocampus (t(35) = 0.96, p = 0.34), nor occipital (t(35) = -0.38, p = 0.71) or parahippocampal (t(35) = 1.57, p = 0.13) cortices (main effect of region: F(2,70) = 1.31, p = 0.28).

Although the hippocampus was the only region that exhibited reliable decoding for Prediction of B, there was no main effect of region. To better understand differences between prediction and perception across regions, we ran two targeted region x comparison ANOVAs. First, we compared Prediction of B and Perception of B, which holds the classifier training data constant (train on B categories, test on A and B trials, respectively). There was no main effect of region (F(2,70) = 2.02, p = 0.14), but there was a reliable main effect of condition (F(1,35) = 7.42, p = 0.010) and a reliable interaction between region and condition (F(2,70) = 7.45, p = 0.0012). Second, we compared Perception of A and Prediction of B, which holds the classifier test data constant (train on A and B categories, respectively, test on A trials). There was again no main effect of region (F(2,70) = 1.23, p = 0.30), but there was a marginal main effect of condition (F(1,35) = 2.96, p = 0.094) and a reliable interaction between region and condition (F(2,70) = 9.85, p *<*0.001). These results suggest a dissociation whereby occipital and parahippocampal cortices more strongly represent perceived information, whereas the hippocampus represents predicted information.

Given that A categories could not be reliably decoded in the hippocampus, we examined whether there was a trade-off in the hippocampus between category evidence for A and B during the viewing of A, with reliable Prediction of B (train on B, test on A) but not Perception of A (train on A, test on A). We assessed this by grouping participants based on whether they exhibited above-chance or chance-level Prediction of B decoding and then comparing Perception of A decoding between these subgroups. We performed this categorical analysis rather than a correlation to account for the fact that we expect variance at or below chance to be noise. We found evidence for a trade-off: participants with above-chance classification for the upcoming B category (vs. other B categories) had lower classification accuracy for the current A category (vs. other A categories) (t(34) = 2.23, p = 0.033). Importantly, we did not find the same relationship between Prediction of B and Perception of X (t(34) = 1.07, p = 0.29). This suggests that prediction can specifically interfere with the ability of the hippocampus to represent the current item.

The analyses above compared the average classification accuracy across participants against an assumed binary chance level of 0.5. Chance classification can deviate from hypothetical levels for a variety of reasons, so we also performed a non-parametric analysis in which we compared classification accuracy against an empirical null distribution estimated for each participant (see Materials and Methods). This analysis yielded nearly identical results (Figure S3).

Given that many regions of the brain have been implicated in predictive processing (Kveraga et al., 2007), we ran an exploratory whole-brain search-light analysis to assess whether regions outside the hippocampus also exhibited Prediction of B decoding (train on B, test on A). No clusters survived correction for multiple comparisons (Figure S4A), consistent with this effect being relatively specific to the hippocampus ROI. To validate the sensitivity of our approach, we repeated the analysis on Perception of B decoding (train on B, test on B). Several regions emerged consistent with our *a priori* ROIs in visual cortex (Figure S4B).

### Relation Between Neural Prediction and Memory Behavior

Finally, we tested our key hypothesis that prediction from statistical learning in the hippocampus is related to impaired encoding of predictive items into episodic memory. We quantified this brain-behavior relationship by correlating (i) each participant’s decoding accuracy for prediction of B during A in the hippocampus with (ii) their difference in hit rate for A vs. X categories in the memory test, which quantifies the relative deficit in memory for predictive items (Figure 3). Consistent with our hypothesis, classification accuracy was negatively correlated with this memory difference (r = -0.33, bootstrap p = 0.047, two-tailed). That is, greater neural evidence for prediction of the upcoming category was associated with worse encoding of the current exemplar.

**Figure 3:**
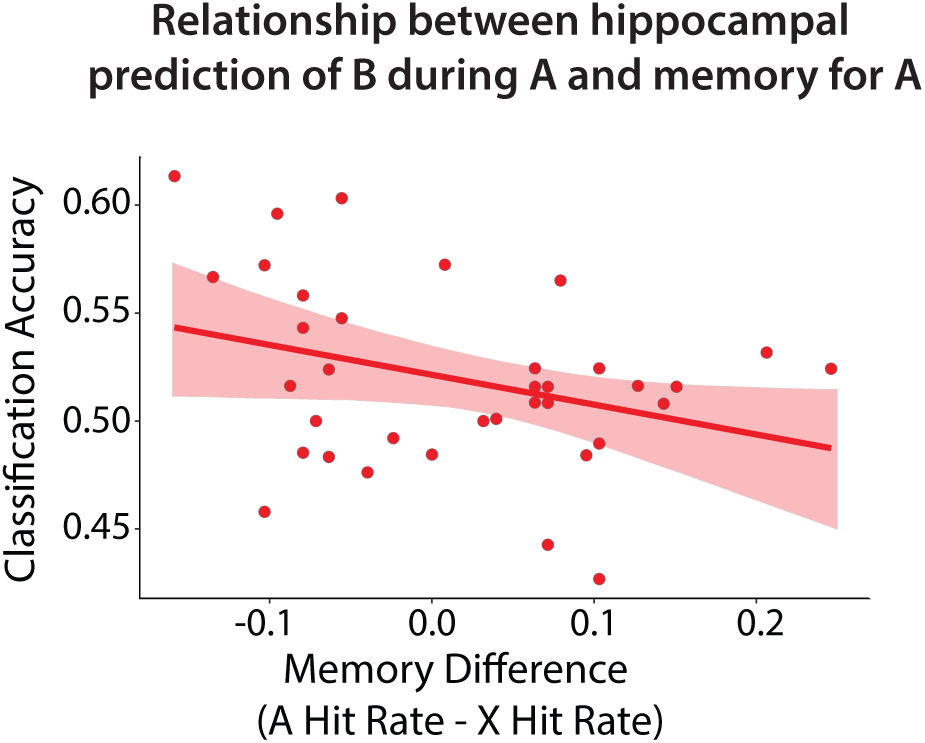
Brain-Behavior Relationship in fMRI Experiment. Pearson correlation between “Prediction of B” classification accuracy in the hippocampus during the encoding phase and the difference in hit rate between A and X. The negative relationship indicates that greater hippocampal prediction from statistical learning was associated with worse episodic memory for the predictive item (see also Figure S5). Error shading indicates bootstrapped 95% confidence intervals.

We included all participants in the correlation above, based on the fact that we observed reliable hippocampal prediction of B at the group level. However, some individuals had decoding accuracy at or below chance, which we do not interpret as meaningful variance. To ensure that these individuals were not driving the negative correlation, we reran the analysis limited to participants with above-chance prediction of B. If anything, the correlation got stronger (Figure S5; r = -0.63; bootstrap p *<*0.001, two-tailed).

## Discussion

Our findings suggest that prediction from statistical learning can interfere with encoding into episodic memory, a process that may be mediated by the hippocampus. Across our behavioral studies (Experiments 1a, 1b, 2), we demonstrated a novel competitive interaction between prediction and memory, such that items which afford an accurate prediction of the upcoming category were remembered worse than items which were unrelated to prediction. In a subsequent fMRI study (Experiment 3), the magnitude of impaired memory for predictive items was associated with neural evidence for the upcoming category in the hippocampus during these items.

### Relation to other studies of the hippocampus

Our findings contribute to growing evidence that the hippocampus plays an important role in statistical learning (Schapiro et al., 2017; Davachi and DuBrow, 2015), including the component processes of prediction (Hindy et al., 2016; Kok and Turk-Browne, 2018) and generalization (Schlichting et al., 2017). In linking these functions to episodic memory, we integrate statistical learning with a broader literature on the role of the hippocampus in memory encoding and retrieval. Specifically, our findings resonate with the observation that encoding and retrieval have fundamentally different requirements (Norman and O’Reilly, 2003; Hunsaker and Kesner, 2013; Neunuebel and Knierim, 2014). Given a partial match between the current experience and past experiences, encoding leverages pattern separation based on the unique features of the current experience and stores a new trace, but in doing so limits access to related old traces. In contrast, retrieval invokes pattern completion to fill out missing features from past experiences and access old traces, but in doing so impedes the storage of a distinct, new trace. To resolve this incompatibility, the hippocampus may toggle between encoding and retrieval states on the timescale of milliseconds to seconds (Hasselmo et al., 2002; Duncan et al., 2012). In the present study, if seeing a picture from an A category triggers pattern completion and activation of its associated B category, the hippocampus may be pushed into a retrieval state that suppresses concurrent memory encoding.

Our findings also resonate with recent findings that the hippocampus represents retrieved information more robustly than perceived information, whereas the visual cortex stably represents both perceived and retrieved information (Lee et al., 2019). These findings are useful to consider in light of the pattern of decoding results that we see across prediction and perception. First, our hippocampal results are consistent with a preference of the hippocampus in representing retrieved information. For example, we found that during the presentation of A, the predicted B information but not the perceived A information could be decoded in the hippocampus. Such a finding could arise from a bias of the hippocampus to represent retrieved information, given a conflict between perception and retrieval. This might also shed light on why only the perception of (predictable) B items, but not A or X items, could be reliably decoded in the hippocampus. Namely, B items may have benefited from retrieval of their category during A. However, this stronger representation of B items in the hippocampus did not translate to consistent evidence for enhanced subsequent memory of B items. This raises intriguing questions about potential differences between prediction and retrieval (Barron et al., 2020).

Second, the pattern of results we find across ROIs are in line with the proposed dissociation between hippocampus and visual cortex in perception and retrieval. Although we did not find strong evidence for a dissociation among ROIs during prediction, the effects were only reliable within the hippocampus. To the extent that the visual cortex is biased toward perceived information and the hippocampus toward retrieved information (Lee et al., 2019), these regions may prioritize different representations during the predictive A items. Whereas the hippocampus may prioritize the predicted B category (as discussed above), and this information may be reinstated in parahippocampal and occipital cortices, the concurrent perceptual information about the A category may dominate and obscure any evidence for B. Indeed, prior studies demonstrating memory-based reinstatement from the hippocampus in visual cortex (Bosch et al., 2014; Tanaka et al., 2014; Hindy et al., 2016; Danker et al., 2017; Kok and Turk-Browne, 2018) were careful to ensure that no conflicting sensory information was present.

Lastly, the findings that the hippocampus preferentially represents retrieved information (Lee et al., 2019) suggest that perceived and retrieved information are coded distinctly in the hippocampus, and thus raise questions about the nature of hippocampal representations during prediction and perception in our task. In particular, the finding that the predicted B category, but not the perceived A category itself could be decoded during A, is especially notable because all of the classifiers were trained on the perception of these categories prior to any learning. Because the categories were counterbalanced across participants, a classifier trained on the perception of a category worked better during A when that category was one of the predicted B categories than when it was one of the perceived A categories. That is, the classifiers generalized from perception to prediction better than from perception to perception, at least in cases where prediction and perception conflicted (i.e., during A). These findings suggest that the format of perceived and predicted information in the hippocampus is similar, consistent with evidence of item-specific reinstatement of perceived information in the hippocampus during retrieval (Mack and Preston, 2016; Tompary et al., 2016).

### Relation to models of learning and memory

How is this interaction between prediction and encoding implemented in the circuitry of the hippocampus? A recent biologically plausible neural network model of the hippocampus (Schapiro et al., 2017) suggests that episodic memory and statistical learning depend on different pathways, the trisynaptic pathway (TSP) and monosynaptic pathway (MSP), respectively. The TSP consists of connections between entorhinal cortex (EC), dentate gyrus (DG), the CA3 sub-field, and the CA1 subfield. In this pathway, DG and CA3 have sparse activity because of high lateral inhibition, which allows them to form distinct representations of similar experiences (i.e., pattern separation) and avoid interference between episodic memories. The MSP consists of a direct connection between EC and CA1. In this pathway, CA1 has lower inhibition and thus higher overall activity and less sparsity, which leads to overlap in the representations of similar experiences, emphasizing their common elements or regularities. Notably, both the TSP and MSP converge on CA1, which is one potential locus of conflict between episodic memory and statistical learning. Future studies tailored for connectivity analysis and/or employing time-resolved and spatially precise methods such as microelectrode intracranial EEG in humans or cellular imaging or recording in animal models are needed to better understand how the hippocampal circuit arbitrates between these two forms of learning.

Importantly, prior models of how the brain processes both episodic memories and statistical regularities have focused on a division of labor between the hippocampus and neocortex, respectively (McClelland et al., 1995). A key distinction between this traditional view and more recent incarnations (Schapiro et al., 2017) is the timescale of statistical learning: the MSP of the hippocampus is well-suited to learning regularities on the order of minutes to hours, whereas the neocortex has a slower learning rate, better suited to extracting regularities over days and weeks. Thus, the neocortex may be important for extracting regularities that span repeated experiences that are spaced out in time in the service of long timescale *semantic* memory, whereas the hippocampus may be preferentially important for regularities experienced repeatedly within the current environment. If true, the hippocampus may consolidate not only discrete episodes into cortex but also these short timescale regularities. The relationship between short and long timescale regularities remains an interesting area for future research. This framework also raises questions about predictions from semantic memory and whether they would interfere with episodic encoding. A key distinction from the kind of rapid statistical learning tasks employed here is that such semantic memory-based predictions may emanate through spreading activation within neocortex, anatomically shielding them from interference by episodic encoding in the hippocampus. However, whether semantic and episodic memories can be fully dissociated remains unclear (Renoult et al., 2019), suggesting a potential interaction of prediction from semantic memory with episodic memory.

In considering the role of the hippocampus in mediating between prediction and encoding, it is important to note that episodic memories can themselves be used to form predictions about the future (Szpunar et al., 2014). This may be particularly useful when regularities are sparsely distributed in time or space and may enable learning via prediction error (Kim et al., 2014) and hypothesis testing (Berens et al., 2018). The extent to which our finding that prediction interferes with episodic encoding reflects a domain-general effect of prediction on memory or is limited to prediction from statistical learning is an exciting question for future research.

### Characterizing the role of prediction in memory

Our work also raises future questions about the nature of the competition between prediction and encoding. After learning predictive relationships in classical conditioning, “blocking” can occur when new cues are introduced. After one conditioned stimulus (CS1) has been paired with an unconditioned stimulus (US), no associative learning occurs when a second conditioned stimulus (CS2) is added (Kamin, 1969). This is interpreted as CS2 being redundant with CS1, that is, not providing additional predictive value given that the US can be fully explained by CS1. In the present study, the A pictures contain two kinds of features: those that are diagnostic of the category (e.g., sand and water for a beach) and those that are idiosyncratic to each exemplar (e.g., particular people, umbrellas, boats, etc.). If categorical features are sufficient to predict the upcoming B category, idiosyncratic features may not be attended or represented (Mackintosh, 1975; Kruschke, 2001), impeding the formation of episodic memories. Our findings are not fully consistent with this account, however. Blocking might predict that the A pictures are represented more categorically, as this is what enables prediction of the B category. Yet, during the presentation of A pictures we found a trade-off in the hippocampus between neural evidence for perception of the A category and prediction of the B category. Nevertheless, more work is needed to better characterize the deficit in memory for predictive items. Are certain aspects of these memories lost while others are retained? Or are these experiences encoded with less precision overall and/or subject to heightened interference at retrieval? Characterizing associative memory between specific A and B exemplars might be a fruitful avenue for future investigation.

### Limitations of the current study

The fMRI study supports and extends the behavioral studies, but has two primary limitations. First, although individual differences in the key memory effect (lower hit rate for predictive vs. control items) were related to neural evidence of hippocampal prediction in Experiment 3, this behavioral effect was not reliable at the group level, as it was in Experiment 1a and replicated in Experiment 1b. Future studies could improve the design to strengthen learning in the scanner environment, including by training the classifier models in a way that does not require pre-learning templates which may have reduced learning (e.g., based on training data from different participants or sessions). Second, a strong version of our hypothesis would suggest that the amount of evidence for the predicted B category on any given A trial should be related to subsequent memory for that specific A item, namely a negative correlation across trials within participant. However, we only found evidence for this relationship by first averaging classifier evidence and subsequent memory within participant and then computing the correlation across participants. Methods that provide cleaner and more time-resolved measurements, such as intracranial EEG, may be better able to resolve neural evidence on single trials in order to examine item-specific relationships.

## Conclusions

Stepping back, why are the computationally opposing functions of episodic memory and statistical learning housed together in the hippocampus? We propose that this shared reliance might allow them to regulate each other. By analogy, using your right foot to operate both the brake and gas pedals in a car serves as an anatomical constraint that forces you to either accelerate or decelerate, but not both at the same time. A similarly adaptive constraint may be present in the hippocampus, reflecting mutual inhibition between episodic memory and statistical learning. When predictive information is available in the environment, further encoding may be redundant with existing knowledge. Moreover, encoding such experiences could risk over-fitting or improperly updating known, predictive regularities with idiosyncratic or noisy details. By focusing on upcoming events, the hippocampus can better compare expectations and inputs (Kumaran and Maguire, 2006), prioritizing the encoding of novel and unexpected events (Greve et al., 2017; Henson and Gagnepain, 2010).

## Author Contributions

Conceptualization, B.E.S. and N.B.T-B.; Methodology, B.E.S. and N.B.T-B.; Investigation, B.E.S.; Formal Analysis, B.E.S.; Writing, B.E.S. and N.B.T-B.; Supervision, N.B.T-B.; Funding Acquisition, B.E.S. and N.B.T-B.

### Acknowledgments

This work was supported by NSF GRFP to B.E.S., as well as NIH R01 MH069456, NSF CCF 1839308, and the Canadian Institute for Advanced Research to N.B.T-B.

## Materials & Methods

### Experiment 1a

#### Participants

Thirty individuals (19 female; age range: 18-31, mean age = 21.2) were recruited from the Yale University community for either course credit or $10 compensation. Informed consent was obtained in a manner approved by the Yale University Human Subjects Committee.

#### Stimuli and Apparatus

Participants were seated approximately 50cm away from a 69cm monitor (1920 × 1080 pixel resolution; 60 Hz refresh rate). Scene stimuli consisted of 300 unique scene images drawn from 12 scene categories (25 images/category), collected from Google image searches. Each participant viewed 22 scenes from each category, randomly selected from the set of 25. Sixteen of these images (per each category; 192 total) were used in the encoding phase, two for the category pair test, and four as foils in the recognition test. Scene stimuli were presented centrally and subtended 27.8 × 20.8 degrees of visual angle. Stimuli were presented using MATLAB (The MathWorks, Natick, MA) with the Psychophysics Toolbox (Brainard and Vision, 1997; Pelli and Vision, 1997).

#### Procedure

Participants first completed an encoding phase. On each trial, they viewed a photograph of a scene for 1000 ms, during which they had to respond based on whether it contained a manmade object (Figure 1A). Participants were instructed to respond as quickly and accurately as possible (response mappings of ‘j’/’k’ onto ‘yes’/’no’ were counterbalanced across participants), and we recorded response time and accuracy. The scene remained on the screen for 1000 ms regardless of button press to equate encoding time, and trials were separated by a 500-ms inter-stimulus interval (ISI) during which a fixation cross appeared.

Every scene was trial-unique, but was drawn from one of 12 outdoor scene categories (beaches, bridges, canyons, deserts, fields, forests, lakes, lighthouses, marshes, mountains, parks, and waterfalls; Figure 1B). Each scene category appeared 16 times over the course of the encoding phase, for a total of 192 trials. The photographs for half of the scene categories always contained a manmade object, and thus all exemplars in a category required the same response, and the responses were balanced overall. Unbeknownst to participants, and orthogonal to the required response, half of the scene categories were assigned to pairs. Given the first scene in a pair (A category scenes), the category of the second scene (B category scenes) was predictable with a transition probability of 1.0. The other half of scene categories were neither predictive nor predictable (X category scenes). Pictures from these categories were inserted on their own randomly, with the constraint that they could not be placed between an A category scene and a B category scene. The assignment of scene categories to A/B/X conditions was itself randomized for each participant. The order of the photograph sequence was randomized with the following three constraints: category pairs and pairs of category pairs could not repeat back-to-back (i.e., no *A*_1_*B*_1_*A*_1_*B*_1_ or *A*_1_*B*_1_*A*_2_*B*_2_*A*_1_*B*_1_*A*_2_*B*_2_, where 1 and 2 index different exemplars); repetitions of each category were spread equally across quartiles of the encoding phase to minimize differences in study-test lag between categories; and the overall transition probability between “yes” and “no” responses on the manmade cover task was forced to be statistically indistinguishable from 0.5.

After the encoding phase, participants performed five minutes of a distracting math phase to minimize recency effects. Each of 60 math problems consisted of division and subtraction, and the answer to the problem was always 1, 2, 3, or 4. Participants responded using the 1, 2, 3, and 4 keys on the keyboard, with a maximum response window of 5 s. The ISI was adjusted based on the response time (5 s minus response time), to ensure that this phase lasted exactly 5 min given the 60 trials. Participants were instructed to respond as accurately as possible.

Participants then underwent two surprise memory tests (category pair test and episodic memory test), the order of which was counterbalanced across participants. The category pair test involved explicit judgments of the category pairings from the encoding phase. Participants were presented with two pairs of photographs on every test trial and were asked to indicate which pair felt more familiar based only what they had seen during the encoding phase. The pairs were shown sequentially: the first scene from one pair appeared for 1000 ms, followed by a 500-ms blank interval, followed by the second scene of the pair for 1000 ms; after a 1000 ms gap with a fixation cross, a second pair was presented in the same manner. After both pairs, participants responded using the ‘1’ key to indicate if the first pair felt more familiar or the ‘2’ key if the second pair felt more familiar. Participants had a maximum of 6 s to respond. Each scene in the category pair test was a completely novel exemplar of its category. Half of the test trials contained a true category pair (when it was a trial testing a pair from the encoding phase); whether it appeared first or second was counterbalanced. The other half of the trials contained a “dummy coded” pair of the X categories (there was no correct answer on these trials). This was done to equate the frequency of categories, which was important for participants who received the category pair test before the episodic memory test. Each true/dummy-coded pair was tested twice against a scrambled pair of the same categories (e.g., if beach → field, mountain → bridge, canyon → forest were category pairs from the encoding phase, the foils might be beach → bridge, mountain → forest, canyon → field). Performance on this category pair test for true pairs vs. scrambled pairs was not reliable in either Experiment 1a (mean accuracy = 0.48; vs. 0.5 chance: t(29) = -0.72, p = 0.48) or Experiment 2 (mean accuracy = 0.49; vs. 0.5 chance: t(29) = -0.61, p = 0.55), nor did the order of the category pair test and episodic memory test affect episodic memory behavior. Thus, the results of the category pair test are not reported further and this test was not included in Experiment 1b and Experiment 3.

The episodic memory test was designed to assess episodic memory for the trial-unique scenes from the encoding phase. On each trial, one scene was presented and participants indicated whether it was “old” (i.e., presented during the encoding phase) or “new” (i.e., not previously seen in the experiment). After making an old/new response (using ‘j’/‘k’ keys on the keyboard), participants then rated their confidence in this response (“not confident”/“confident”, using ‘d’/‘f’ keys). Participants had 6 s to make each response. All 192 scene photographs from the encoding phase were shown, in addition to 48 foils (4 novel exemplars from each category). The order of the scenes was randomized.

### Experiment 1b

#### Participants

Eighty-three individuals were recruited from the online data collection platform Prolific. All participants self-reported that they were between the ages of 18 and 35, had normal or corrected-to-normal vision, and lived in the U.S. or the U.K. Informed consent was obtained in a manner approved by the Yale University Human Subjects Committee. Nineteen participants were excluded based on pre-registered criteria, resulting in 64 usable participants, in line with our pre-registered sample size. The full pre-registration for this study can be found here: https://aspredicted.org/blind.php?x=my8ky2.

#### Stimuli and Apparatus

Scene stimuli were the same as in Experiment 1a. Stimuli were presented using custom Javascript code for online testing.

#### Procedure

The procedure was identical to Experiment 1a, except for the following changes: during the encoding phase, all participants responded with ‘j’ key for yes and ‘k’ key for no; the math distractor task was simplified to contain only subtraction problems (no division); and no category pair test was administered.

### Experiment 2

#### Participants

Thirty individuals (19 female; age range: 18-23, mean age = 19.3) were recruited from the Yale University community for either course credit or $10 compensation. Informed consent was obtained in a manner approved by the Yale University Human Subjects Committee.

### Stimuli and Apparatus. Same as Experiment 1a

#### Procedure

The procedure was identical to that of Experiment 1a, with the addition of a temporal source memory judgment in the test phase. That is, participants were presented with a scene and first asked to judge whether it was “old” or “new” (using the ‘d’ and ‘f’ keys). Then, old responses were followed by the presentation of a timeline, bound by the start and end clock times of the encoding phase. Participants used the mouse to click along the timeline to indicate when they remembered seeing the scene. No temporal source judgments were collected after new responses.

### Experiment 3

#### Participants

Thirty-eight individuals (24 female; age range: 18-35, mean age = 23.1) were recruited from the Yale University community for $30 compensation. We chose this slightly higher sample size than the in-person behavioral studies (Experiments 1a and 2) under the assumption of a 20% attrition rate in MRI studies, aiming for the same sample size of 30 participants. One participant was excluded to due a neurological anomaly and one participant was excluded for chance-level episodic memory performance (overall A′ *<*0.5). Additionally, one participant was excluded from response time analyses because of a technical error that resulted in no responses being collected for part of the encoding phase. Thus, the final sample size for fMRI analysis and memory performance was 36 participants. Informed consent was obtained in a manner approved by the Yale University Human Investigation Committee.

#### Stimuli and Apparatus

Stimuli were presented on a rear-projection screen using a projector (1920 × 1068 pixel resolution; 60 Hz refresh rate). Participants viewed the stimuli through a mirror mounted on the head coil. Scene stimuli consisted of 480 unique images drawn from 12 categories (40 images/category), collected from Google image searches (180 additional stimuli were collected for this experiment; the other 300 are identical to those used in Experiments 1 & 2). Each participant viewed 39 scenes from every category, randomly selected from the set of 40: 21 per category (252 total) for the encoding phase, 14 (168) for the pre and post templating phases, and 4 (48) as foils in the episodic memory test.

#### Procedure

The procedure was identical to Experiment 1a other than the following changes:

Instead of one continuous block of the encoding phase (with 16 repetitions of each scene category), the stream was divided into three fMRI runs, each with seven repetitions/category (such that were 21 repetitions/category in total across the encoding phase). As in Experiments 1a, 1b, and 2, each image was presented for 1 s, but the ISI varied between 2 s (39.3% of trials), 3.5 s (39.3% of trials), and 5 s (21.4% of trials) to jitter onsets for deconvolving event-related fMRI activity. For the manmade object cover task, participants responded using their right index and middle fingers on an MR-compatible button box.

Before and after the three runs of the encoding phase there were “pre” and “post” templating phases (one fMRI run each). To participants, these phases were identical to the encoding phase (e.g., stimulus timing and task were identical). However, there were no category-level regularities in these two runs. Scenes from all categories were presented in a random order. To limit the impact of this random presentation on subsequent learning, participants completed a distracting math task between the “pre” templating run and the first encoding phase run. Each of these five functional runs (three encoding phase runs and pre/post runs) lasted 6.4 minutes.

For the episodic memory test, as in Experiment 1a and 1b, a scene was presented and participants indicated (with their index and pinky fingers) whether it was “old” (i.e., presented during the encoding phase) or “new” (i.e., not previously seen in the experiment). They then rated their confidence in this response (“very unsure”,“unsure”,“sure”,“very sure”), using their index through pinky fingers, respectively. Participants had 6 s to respond to each of these questions. They completed this task while in the scanner, but no fMRI data were collected. No category pair test was administered in this experiment. *MRI Acquisition*. Data were acquired on a Siemens Prisma 3T scanner using a 64-channel head coil at the Magnetic Resonance Research Center at Yale University. Functional images were acquired using an EPI sequence with the following parameters: TR = 1500 ms; TE = 32 ms; 90 axial slices; voxel size = 1.5 × 1.5 × 1.5 mm; flip angle = 64 degrees; multiband factor = 6. Additionally, a pair of opposite phase-encode spin-echo volumes were collected for distortion correction (TR = 11,220 ms; TE = 66 ms). One T1-weighted MPRAGE (TR= 1800 ms; TE = 2.26 ms; voxel size = 1 × 1 x 1 mm; 208 sagittal slices; flip angle = 8 degrees) and two T2-weighted turbo spin echo (TR = 11,390 ms; TE = 90 ms; 54 coronal slices; voxel size = 0.44 × 0.44 × 1.5 mm; distance factor = 20%; flip angle = 150 degrees) anatomical images were collected.

#### fMRI Preprocessing

fMRI data processing was carried out using FEAT (fMRI Expert Analysis Tool) Version 6.00, part of FSL (FMRIB’s Software Library, www.fmrib.ox.ac.uk/fsl) version 5.0.10. EPI and anatomical images were skull-stripped using the Brain Extraction Tool (Smith, 2002). Susceptibility-induced distortions measured via the opposing-phase spin echo volumes were corrected using FSL’s topup tool (Andersson et al., 2003). Each functional run was high-pass filtered with a 128 s period cut-off, corrected for head motion using MCFLIRT (Jenkinson et al., 2002), and motion outliers were computed. Slice-timing correction was performed. No spatial smoothing was applied. Lastly, the six motion parameters, as well as motion outliers, were regressed against the BOLD timecourse using a general linear model (GLM). The residuals from this preprocessing model (which contain BOLD responses to the task after controlling for motion) were then used for subsequent analyses.

Functional images were registered to each participant’s T1 anatomical scan using boundary-based registration, as well as to a 2 mm MNI standard brain, using 12 degrees of freedom. Lastly, the two T2 anatomical images collected were registered to one another and averaged; the resulting averaged image was registered to the T1 anatomical image using FLIRT (Jenkinson and Smith, 2001).

#### Regions of Interest

The hippocampus region of interest (ROI) was defined anatomically by concatenating subfields CA1, CA2/3/DG, and subiculum. These hippocampal subfields, as well as surrounding medial temporal lobe (MTL) cortical regions including the parahippocampal cortex (PHC) ROI, were manually segmented on each participant’s averaged T2 anatomical scan using published anatomical landmarks (Insausti et al., 1998; Pruessner et al., 2002; Duvernoy, 2005; Aly and Turk-Browne, 2015). The occipital cortex ROI was defined using the MNI structural atlas, thresholded at 25% probability. For each region, we concatenated across left and right hemispheres to create one bilateral ROI, as we had no hemisphere-specific hypotheses. These ROIs were then transformed into the participant’s functional space for subsequent analyses.

#### Category Decoding Analysis

A multivariate pattern classification approach was used to assess evidence for a particular category during the encoding phase. This approach involved training a classifier on fMRI activity patterns for each category from the “pre” templating run (when there were no regularities present and none had been learned) and testing for classifier evidence of these categories during the three (independent) runs of the encoding phase. We tested for category evidence during all three runs of the encoding phase, given prior work demonstrating that neural evidence of statistical learning can occur quite rapidly (Turk-Browne et al., 2009, 2010). Although the beginning of the first run may contribute noise (as no learning can possibly have occurred), we erred on the side of including more data.

For each functional run, the residuals from the preprocessing GLM (with known noise sources removed but still containing task responses) were aligned to the final functional run and z-scored across time. The voxel x time matrices were then masked to only include voxels within an ROI. The timepoints corresponding to the presentation of each of two categories of interest were extracted and shifted by 3 TRs (4.5 seconds) to account for the hemodynamic lag. The voxel activity patterns from these shifted time points were then used as training or test data for the classifier. Timepoints that included a motion outlier were excluded from the training/test sets.

Linear SVMs were trained on data and labels from the pre-learning templating run, using the SVC function in Python’s scikit-learn module, with a penalty parameter of 1.00. Classifiers were then tested with data corresponding to the timepoints of the trained categories in the three runs of the encoding phase (concatenated) and made guesses as to the category label of each test example. Accuracy was computed as the proportion of correct guesses.

We ran the following comparisons: Perception of A (training on pre-learning examples of A, testing for evidence of A during the presentation of A in the encoding phase), Perception of B (training on pre-learning examples of B, testing for evidence of B during the presentation of B in the encoding phase), Lingering of A (training on pre-learning examples of A, testing for evidence of A during the presentation of B in the encoding phase), Prediction of B (training on pre-learning examples of B, testing for evidence of B during the presentation of A in the encoding phase), and Perception of X (training on pre-learning examples of X, testing for evidence of X during the presentation of X in the encoding phase).

Each participant encountered three A and three B categories over the course of the experiment. Thus, for each of the four comparisons above, we built three different binary classifiers and then averaged their accuracy. In other words, a classifier was trained to distinguish between two scene categories from the same condition (e.g., B) based on the pre-learning templating run, and tested for evidence of those two categories during the subsequent presentation of two categories (e.g., their corresponding As for Prediction of B) in the encoding phase. Accuracy (percent correct) was then computed for each of these three classifiers (A1 vs. A2; A2 vs. A3; A1 vs. A3) and averaged, resulting in one mean accuracy value per comparison, per participant.

To provide an example, if the category pairs were beach → field, mountain → bridge, canyon → forest, then B classifiers would be trained for field vs. bridge, bridge vs. forest, and field vs. forest. To calculate evidence for Prediction of B: the field vs. bridge classifier would be applied to the beach and mountain trials — such that the classifier estimated evidence for field and bridge during each beach or mountain trial — and accuracy was computed (such that, for example, accuracy on a beach trial was 1 if the classifier outputted more evidence for field than for bridge). This was repeated for the bridge vs. forest classifier (testing for evidence of these categories during mountains and canyons) and the field vs. forest classifier (testing for evidence of these categories during beaches and canyons). The accuracies of these three classifiers were averaged into a single accuracy for each participant. This was repeated for the three other comparisons above and for each ROI. To assess reliability at the group level, performance was compared to a chance level of 0.50 across participants using a one-sample t-test.

To quantify classification accuracy non-parametrically, we performed randomization tests in which we computed an empirical null distribution of classification accuracy values for each participant. The null distributions were generated from 1,000 iterations of shuffling the category labels at test prior to scoring the model. We then calculated a z-score for each participant’s true classification accuracy relative to their own null distribution. To test reliability, we compared these z-scores against 0 across participants (Figure S3).

#### Assessing Reliability of Correlations

To estimate correlations across participants robustly (e.g., as used in the Relation Between Neural Prediction and Memory Behavior section), we performed a random-effects bootstrap resampling procedure (Efron and Tibshirani, 1986). For each of 10,000 iterations, we randomly drew 36 participants from our sample with replacement, and recalculated the Pearson correlation between the two variables of interest. This procedure operates under the assumption that if the effect is reliable across participants, then the participants are interchangeable and which subset is resampled in any given iteration will not affect the outcome. This approach also helps mitigate the impact of outliers when calculating correlations from modest sample sizes. The resulting sampling distribution can be used to generate confidence intervals and perform null hypothesis testing. Specifically, we calculated the p value as the proportion of iterations in which the correlation value was of the opposite sign from the true correlation, then multiplied by 2 for a two-tailed significance.

## Data Availability

fMRI data can be downloaded from OpenNeuro and behavioral data can be downloaded from Dryad.

