## Supplementary material for "Statistical prediction of the future impairs episodic encoding of the present": Figures S1-S5, Table S1

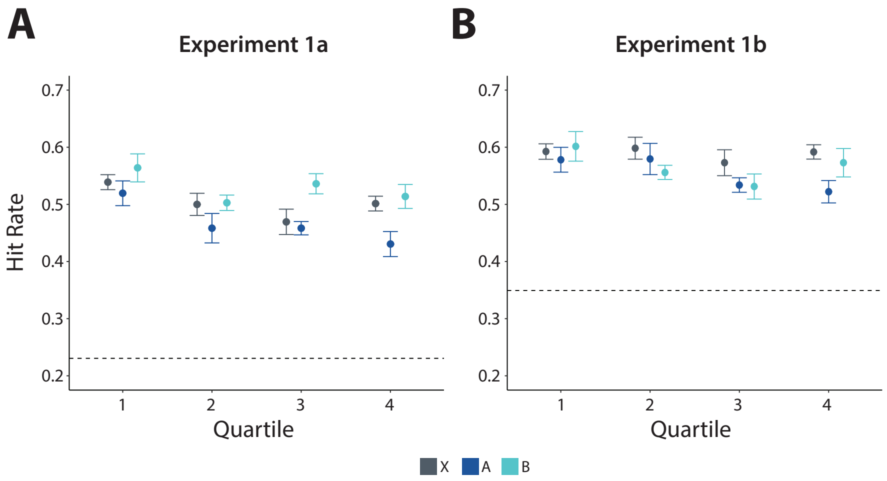


Figure 1: Hit Rate as a function of condition and quartile for (A) Experiment 1a and (B) Experiment 1b. The dashed line indicates the mean false alarm rate.

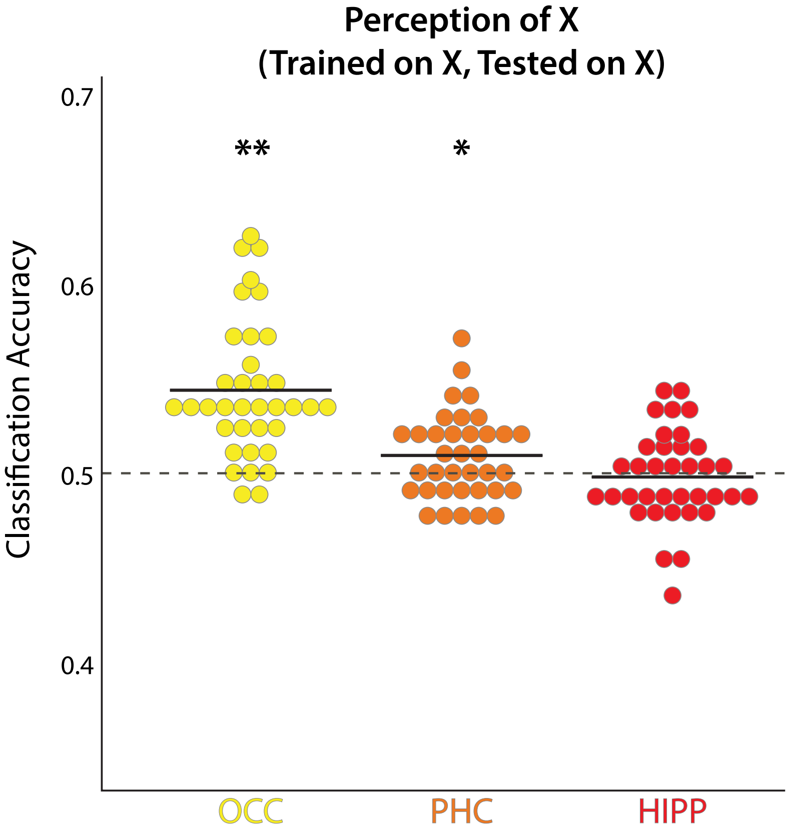


Figure 2: Decoding of non-predictive, non-predictable control X categories in the occipital cortex, parahippocampal cortex, and hippocampus. For each ROI, each dot is one participant and the black line is the mean across participants. OCC: $t(35) = 7.16$, $p < 0.001$; PHC: $t(35) = 2.47$, $p = 0.019$; HIPP: $t(35) = -0.49$, $p = 0.63$. *p <0.05; **p <0.01

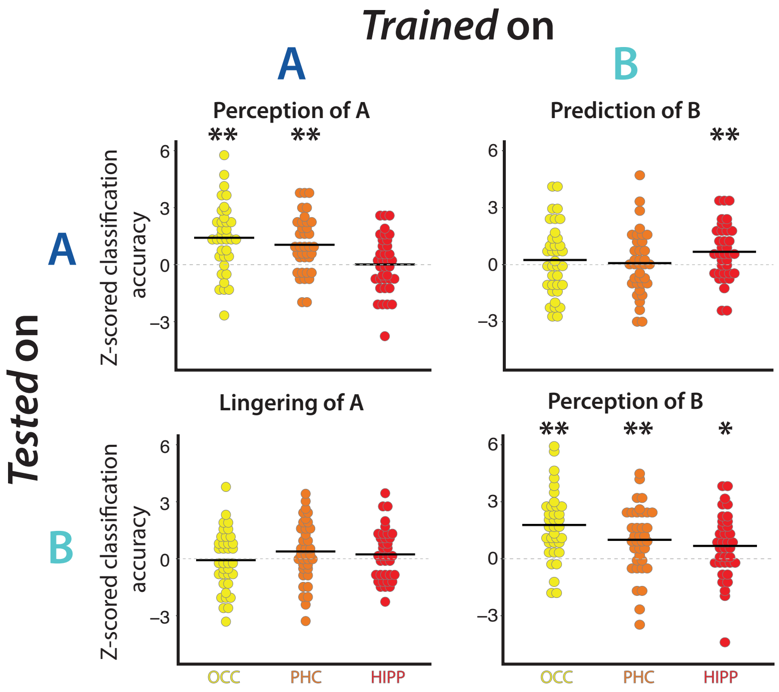


Figure 3: Category Decoding in fMRI Experiment. Z-scored classification in occipital cortex (OCC), parahippocampal cortex (PHC), and hippocampus (HIPP) plotted for each of the four combinations of training or testing on A or B categories. Z-scores were computed by generating a permuted null distribution of classification accuracy values for each participant and calculating the z-score of each participant's empirical classification accuracy relative to their own null distribution. For every A/B combination and ROI, each dot is one participant and the black line is the mean across participants. Perception of A: OCC: $t(35) = 4.65$, $p < 0.001$; PHC: $t(35) = 4.18$, $p < 0.001$; HIPP: $t(35) = 0.043$, $p = 0.97$. Prediction of B: OCC: $t(35) = 0.79$, $p = 0.43$; PHC: $t(35) = 0.29$, $p = 0.77$; HIPP: $t(35) = 2.79$, $p = 0.0084$. Lingering of A: OCC: $t(35) = -0.29$, $p = 0.77$; PHC: $t(35) = 1.43$, $p = 0.16$; HIPP: $t(35) = 1.00$, $p = 0.33$. Perception of B: OCC: $t(35) = 5.89$, $p < 0.001$; PHC: $t(35) = 3.38$, $p = 0.0018$; HIPP: $t(35) = 2.36$, $p = 0.024$. *$p < 0.05$; **$p < 0.01$

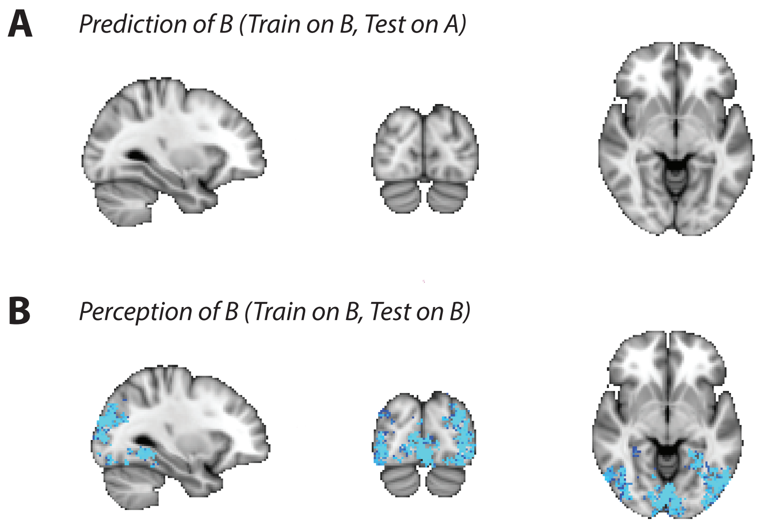


Figure 4: Searchlight results. We conducted a voxelwise analysis to explore the specificity of our Prediction of B results in the brain. We repeated the category decoding analysis in 27-voxel cubes centered on all functional voxels (searchlight function in BrainIAK) (Kumar et al., 2020). Each aligned, normalized residual volume from the pre-processing GLM was registered to standard space. These volumes were masked for each searchlight cube and the retained voxels were subjected to the same decoding pipeline described in the main text for the ROIs. The result was a searchlight map per participant, in which the value at each voxel reflected the average classification accuracy for the cube centered at that voxel. The reliability of these maps was assessed at the group level using non-parametric randomization tests (randomise function in FSL) (Winkler et al., 2014), corrected for multiple comparisons using threshold-free cluster enhancement (Smith and Nichols, 2009). A) No clusters exhibited reliable decoding for Prediction of B (corrected $p < 0.005$). B) As a control analysis, we ran the same searchlight procedure for Perception of B and obtained reliable decoding throughout visual cortex, including occipital and parahippocampal ROIs (corrected $p < 0.005$). Statistical maps are plotted on the T1 MNI standard brain (2mm), at slices X = -30 (sagittal), Y = -82 (coronal), Z = -12 (axial).

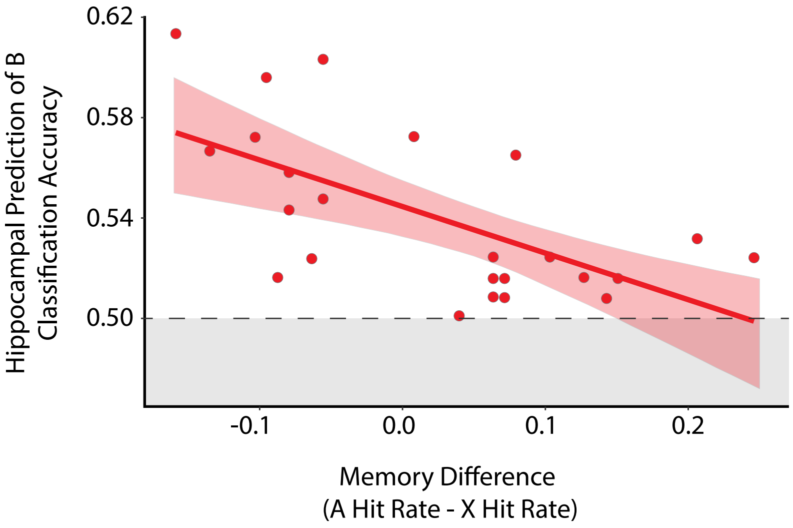


Figure 5: Relation between hippocampal "Prediction of B" classification accuracy and difference in hit rate between A and X, excluding participants with classification accuracy at or below chance (<0.5), marked by gray shading. The negative correlation holds when excluding these participants, showing that they were not driving the overall effect. Error shading indicates bootstrapped 95% confidence intervals.

| Experiment 1a | A | B | X |
|---|---|---|---|
| $A'$ | .700 (.11) | .749 (.089) | .713 (.091) |
| Hit Rate | .467 (.12) | .529 (.11) | .502 (.12) |
| False Alarm Rate | .228 (.14) | .217 (.15) | .239 (.11) |
| **Experiment 1b** | A | B | X |
| $A'$ | .670 (.13) | .672 (.13) | .698 (.088) |
| Hit Rate | .553 (.16) | .565 (.16) | .589 (.14) |
| False Alarm Rate | .348 (.22) | .354 (.21) | .348 (.18) |
| **Experiment 2** | A | B | X |
| $A'$ | .737 (.14) | .737 (.085) | .758 (.073) |
| Hit Rate | .400 (.14) | .369 (.15) | .379 (.14) |
| False Alarm Rate | .125 (.13) | .119 (.097) | .107 (.091) |
| **Experiment 3** | A | B | X |
| $A'$ | .656 (.13) | .697 (.10) | .678 (.079) |
| Hit Rate | .609 (.13) | .633 (.13) | .594 (.14) |
| False Alarm Rate | .412 (.22) | .382 (.19) | .382 (.16) |

Table 1: Average (SD) memory performance in the behavioral test of each experiment.
